# Drivers of plant traits and forest functional composition in coastal plant communities of the Atlantic Forest

**DOI:** 10.1101/812339

**Authors:** Jehová Lourenço, Erica A. Newman, Jose A. Ventura, Camilla Rozindo Dias Milanez, Luciana Dias Thomaz, Douglas Tinoco Wandekoken, Brian J. Enquist

## Abstract

The severe deforestation of Brazil’s Atlantic Forest and increasing effects of climate change underscore the need to understand how tree species respond to climate and soil drivers. We studied 42 plots of coastal *restinga* forest, which is highly diverse and spans strong environmental gradients. We determined the forest physiognomy and functional composition, which are physical properties of a community that respond to climate and soil properties, to elucidate which factors drive community-level traits. To identify the most important environmental drivers of coastal Atlantic forest functional composition, we performed a forest inventory of all plants of diameter 5 cm and above. We collected wood samples and leaves from ∼85% of the most abundant plant species and estimated height, aboveground biomass (AGB), and basal area of individual plants, and the community-weighted specific leaf area (SLA). In addition to plant traits, we measured water table depth and 25 physicochemical soil parameters. We then parameterized several models for different hypotheses relating the roles of nutrients and soil to tropical forest diversity and functioning, as represented by plant traits. Hypotheses were formalized via generalized additive models and piecewise structural equation models. Water table depth, soil coarseness, potential acidity, sodium saturation index (SSI) and aluminum concentration were all components of the best models for AGB, height, basal area, and trait composition. Among the 25 environmental parameters measured, those related to water availability (water table depth and coarse sand), followed by potential acidity, SSI, and aluminum consistently emerged as the most important drivers of forest physiognomy and functional composition. Increases in water table depth, coarse sand, and soil concentration of aluminum negatively impacted all the measured functional traits, whereas SSI had a positive effect on AGB and plant height. These results suggest that sodium is not merely tolerated by Atlantic Forest *restinga* plant communities, but is important to their structure and functioning. Presence of aluminum in the soil had a complex relationship to overall basal area, possibly mediated by soil organic matter.

## Introduction

Tropical tree species’ distributions are fundamentally affected by soil nutrient availability, highlighting and the role of soils in functional assembly and biogeography (Chadwick and Asner 2018), and the importance of resource-competition as a mechanism structuring tropical plant communities (John et al. 2007). Soil plays a fundamental role in the diversity and functioning of tropical forests, and underlies a complex interacting system of forest taxonomic composition and soil formation, driving patterns of niche differentiation at the global scale (Fujii et al. 2018). Soil nutrients, structure, and the water affect tropical forest diversity and function at broad scales (Cole 1960, Webb and Peart 2000, Phillips et al. 2009), driving differences between characteristics of major biomes (such as forest and savanna). Few studies have looked at how these same properties affect local-scale heterogeneity in vegetation physiognomy (de Assis et al. 2011). It remains an open question as to how changes in soil nutrients, texture and chemistry affect local forest structure and physiognomy, which is defined as “the form and function of vegetation; the appearance of vegetation that results from the life-forms of the predominant plants” (Shimwell 1984) (attributed to Cain and Castro, 1959). Studies that focus on strong environmental gradients can help guide our understanding of determinants of ecosystem structural composition, and eventually, ecological responses to environmental changes (Cornwell and Ackerly 2009, Violle et al. 2011, Guittar et al. 2016).

In Brazil’s Atlantic Forest, predictors of coastal *restinga* physiognomy have been attributed to several environmental variables, among which are soil nutrients (Lourenço and Cuzzuol 2009), salinity (Lourenço et al. 2013), organic matter, aluminum saturation (Rodrigues et al. 2013), soil texture and water table depth (Cooper et al. 2017). It has been suggested that water table depth and microtopography in *restinga* forest structure may be the most important variable driving plant traits, because they drive fundamental changes in physicochemical soil properties, such as nutrient content, base saturation, acidity, cation exchange capacity, organic matter, and aluminum saturation (de Almeida et al. 2011), which may constrain *restinga* plant growth (Marques et al. 2015). Such conditions may exert a strong environmental filtering over the forest taxonomic composition, selecting species that are adapted to grow in soils poor in nutrients, and that have high aluminum tolerances (Britez et al. 2002b, 2002a).

Furthermore, the porous *restinga* soils result in low soil fertility and low water retention capacity (Cooper et al. 2017), which may impose constraints on plant biomass productivity (Lambers et al. 2008, Santiago-garcía et al. 2019). In water limited conditions, plants exhibit more conservative strategies, such as small body size and low specific leaf area (Cornwell and Ackerly 2009, Katabuchi et al. 2012), and the production of thick and scleromorphic leaves with increased longevity (Wright et al. 2004, Lambers et al. 2008), as are typically found in *restinga* plant species (Mantuano et al. 2006, De Aguiar-Dias et al. 2012, Melo Júnior and Torres Boeger 2015, Pinedo et al. 2016).

The role of sodium in the Amazon and other tropical forests may be much more general and important than previously thought (Kaspari et al. 2009). Increasing soil salt (NaCl) concentration toward the coast and the influence of the saline spray by the proximity to the ocean (Griffiths 2006) is likely to be an important driver of species distribution in the Atlantic Forest ecosystem. Salt concentration in the soil may act as a filter on the functional and taxonomic composition of coastal plant communities, selecting species with salt tolerant traits, such as high leaf mass per area, thicker leaves, and the capacity for salt exclusion (Poorter et al. 2009). In the other hand, the presence of salt may also accelerate the biological decomposition of the litter in tropical forests (Kaspari et al. 2009), which may have positive consequences to the nutrient cycling of systems poorer in nutrients, such as *restinga*. Moreover, salinity was recently reported as an important predictor of *restinga* forest locations in comparison to other Atlantic Forest marginal habitats (Neves et al. 2017), and we therefore expect that NaCl may be playing a key role in this ecosystem.

Aluminum also plays a potentially large role in determining local physiognomic characteristics, including plant height, aboveground biomass, and other aspects of vegetation form. Aluminum is moderately toxic to a wide range of plants, and it has been shown to facilitate the immobilization of phosphorus. Aluminum may also interfere with plants’ uptake of basic cationic nutrients (Richards 1952), although this relationship changes in acidic soils and may actually facilitate the uptake of nitrogen, phosphorus and potassium in plants adapted low pH soils (Osaki et al. 1997). Elsewhere in the Amazon Basin, potassium has been identified as playing a key role in regulating forest structure (Lloyd 2015), whereas soil clay content, acidity, and moisture saturation levels seem more important in the *Cerrado*, where aluminum is present in significant amounts, but cannot explain local physiognomic heterogeneity (de Assis et al. 2011).

Given the high species richness, extremely high level of endemism and the small fraction of the original forest cover remaining, there has been an increasing need for preservation of Atlantic Forest habitats (Scarano 2009, Joly et al. 2014), which are ranked among the top five hotspots of biodiversity in the world (Myers et al. 2000). The scarcity of studies addressing *restinga* plant-soil relationship highlights the necessity for a more detailed study on the drivers of local forest physiognomy and functional composition, which may be applied in the context of protection and restoration of these highly diverse and threatened plant communities. Moreover, we argue that assessing such drivers underlying *restinga* forest structure may shed some light to broader questions regarding to the factors controlling tropical forest diversity and function.

We address the following questions: 1) what are the main environmental drivers of functional composition in *restinga* forests? and 2) how do the most important environmental drivers influence *restinga* forest physiognomy and functional composition? Through measurement of a suite of plant traits and 25 environmental variables in 42 *restinga* plant communities, we develop a mechanistic understanding of the system functioning, and the interaction between environmental variables and its effects on forest physiognomy and trait composition.

## Material and Methods

### Study area and aboveground surveys

The *restinga* forest is one of the most environmentally and biologically diverse habitats of Atlantic Forest (Thomas and Garrison 1998, Scarano 2009). This environmental diversity is attributed to the conspicuous soils mosaic and topographic variation arising from the recent transgressive and regressive events from the Quaternary (Suguio and Martin 1990). The region is characterized by a conspicuous soil mosaic and a waved relief, which create microenvironmental conditions for the occurrence of steep flooding gradients. The topography affects the water flow which tends to accumulate in lower sites with organic soils, and shallow water table depth (floodable or permanently flooded forests), while upper sites (non-floodable areas) are comprised of well-drained sandy soils, creating continuous soil gradients of water and nutrients availability.

Data collection for this study took place in Paulo Cezar Vinha State Park, which is located in Guarapari municipality of Espirito Santo State, Brazil, and is a *restinga* protected area (Fig. 1), with several discrete plant communities that are found side-by-side. Pereira (1990) reported 11 well-defined plant communities in this park, ranging from small shrubs to tree communities, which show increasing structural complexity from the shore towards the inland areas (Pereira 1990, Assis et al. 2004). The study area (Fig. 1B) covers a short flooding gradient transition, where we set up 42 plots (measuring 5 x 25 meters) across three forest types: floodable, intermediate, and dry forests (Fig. 1C). All trees with diameter at the breast high (dbh) ≥5cm were tagged, and height and dbh were measured. The close spatial scale of communities allowed us to precisely track the strong continuous flooding gradient (207.4 ± 60.7 meters), increasing the sampling accuracy in detecting the effect of soil gradients in wood and leaf traits, and overall forest composition.

**Figure 1.**
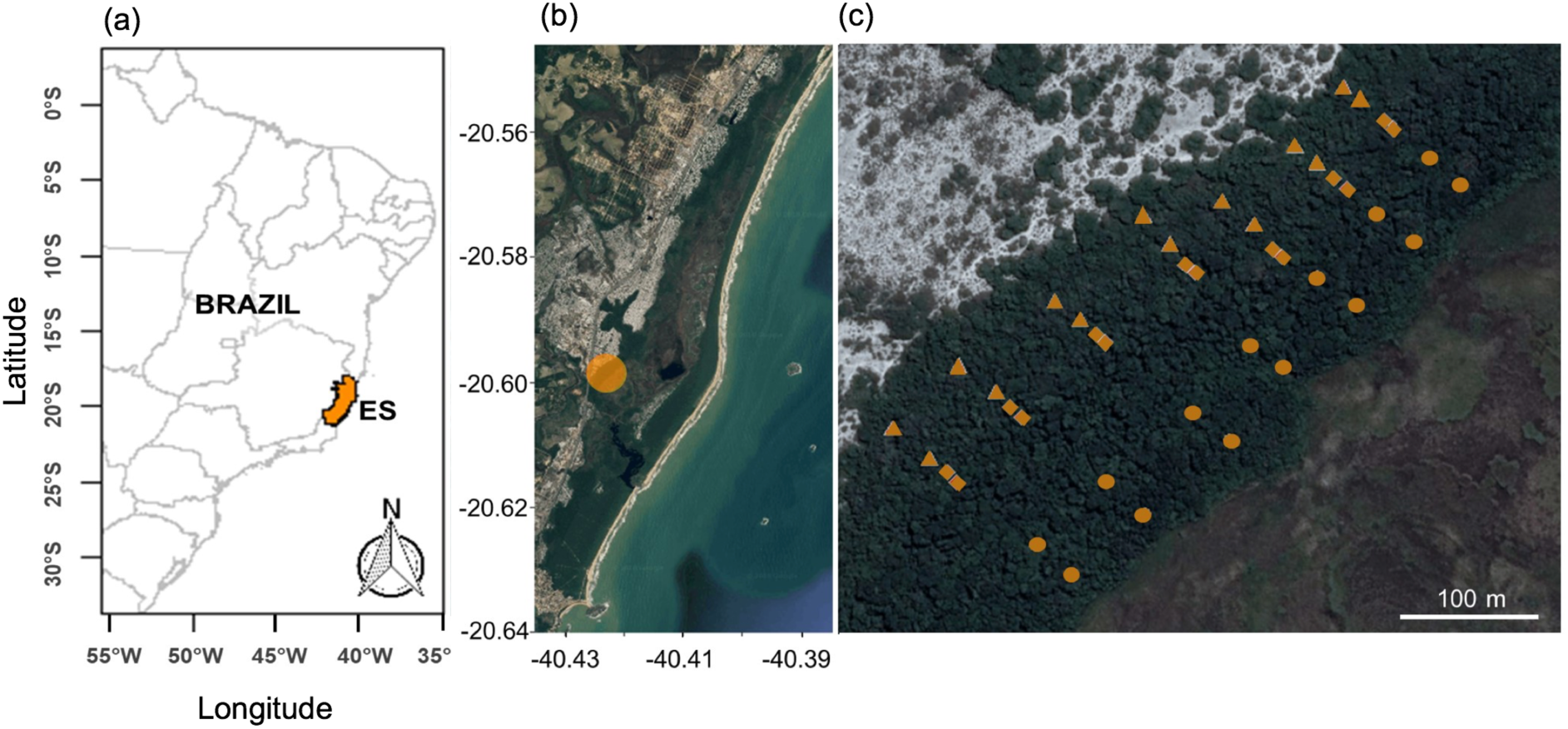
(a) Paulo Cesar Vinha State Park and study area location (orange dot). (b) Detailed map of the study area, showing the 42 plots (or local communities) distributed across floodable (○), intermediate (□), and dry (Δ) sites.

### Measuring and estimating plant traits

We estimated total aboveground biomass of each individual tree with Chave’s pantropical models using the computeAGB function provided by the “BIOMASS” R package (Chave et al. 2014), which requires height, dbh and wood density (WD) data. We collected wood samples from five individuals per species and per site (floodable, intermediate and dry) to accurately determine the wood density of each species in each specific environment. Around 85% of the most abundant plant species were sampled. The samples were then hydrated for 12 hours, and the wood volume was calculated according to the water displacement method (Chave 2005), with the support of a high precision balance. We then dried the samples in an oven at 60°C and weighed them. WD (in g/cm^3^) was determined by dividing the wood dry mass by its volume.

We determined the WD of the 15% less abundant plant species by using a hierarchical approach, which used (as a reference) the mean WD value for the genus or family related species, whose occurrence matched the same site or environmental condition. If the species had no similar genus or family in the study area, the WD was estimated from the getWoodDensity function from R “BIOMASS” package (Chave et al. 2014), using “SouthAmericaTrop” as the “region” argument. For simplicity, each plot was defined as a local plant community. The maps were drawn using the “maptools” (Bivand and Lewin-Koh 2017) and “raster” packages (Hijmans 2017) in the R Statistical Environment (R Core Team, 2018).

### Soil sampling and analysis

Nutritional and physicochemical soil composition were measured in each plot via soil sampling. Five soil samples were collected per plot at a depth of 15 cm, and were then homogenized in the field to produce one compound soil sample per plot. Several parameters were determined by the soil analysis, including coarseness (proportion of fine and coarse sand, silt, and clay); nutrients (P, K, Na, Ca, Mg, Al, H+Al [potential acidity], Zn, Mn, Cu and B); organic matter (OM); pH; sodium saturation index (SSI), aluminum saturation index (ASI), base saturation (BS), base saturation index (BSI), effective cation exchange capacity (CEC) and cation exchange capacity in a pH of 7 (CEC7), following the Brazilian Agricultural Research Corporation protocol (Donagema et al. 2011) (Table 1).

**Table 1.**
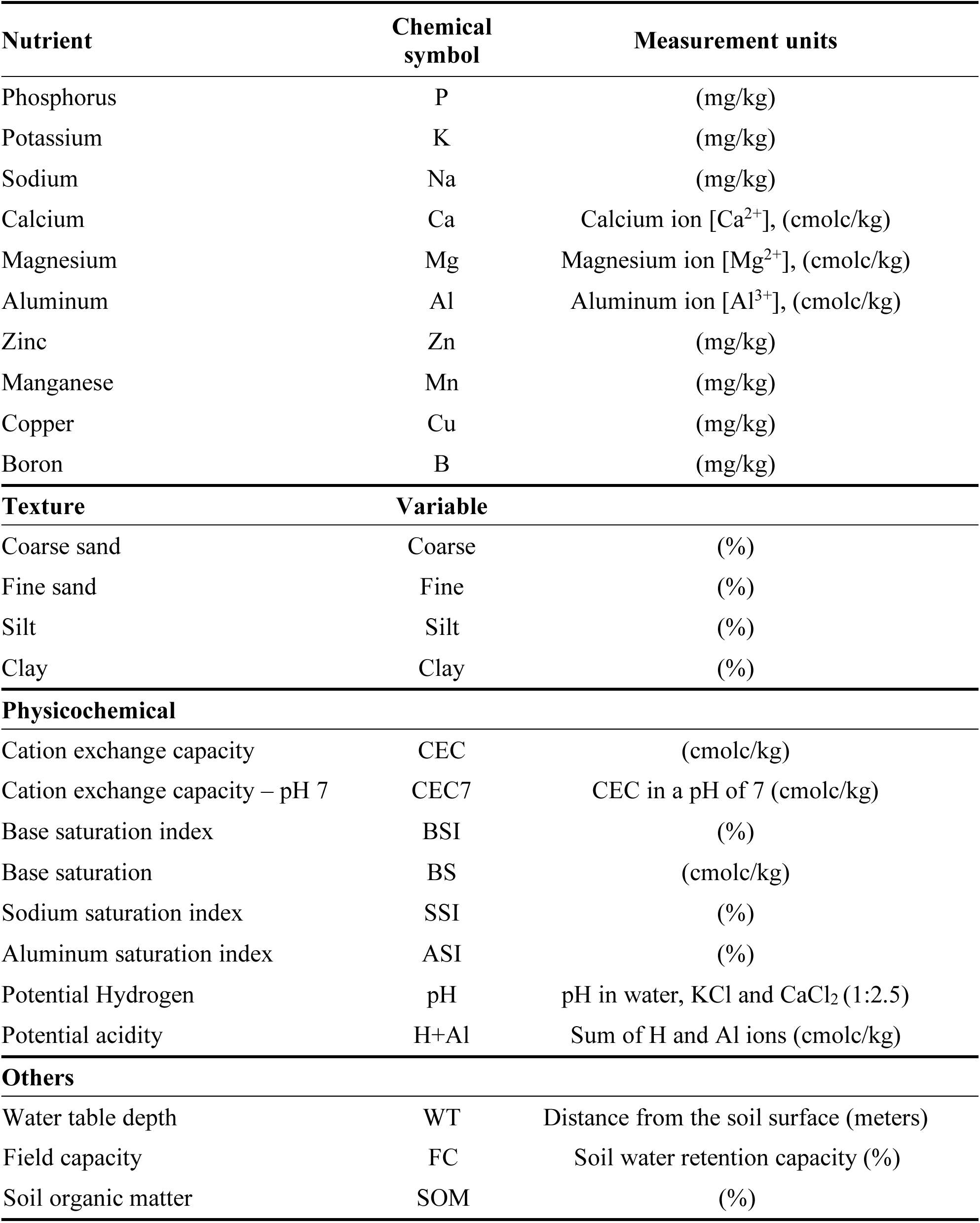
Environmental variables used in forward selection analysis, including nutrients, texture, and physicochemical soil parameters.

We also measured the soil water retention capacity or field capacity (FC) for each plot. This was achieved by collecting two soil samples (15 cm depth) per plot using flexible pipes to remove the soil layer in such a way as to preserve the integrity of their original structure. We introduced water to both samples until the wetted soil samples exceeded their maximum water retention capacity. When water stopped draining from samples, they were weighed in a balance, dried in an oven at 60°C, and daily weighted until they reached a constant weight. The FC was determined following equation: FC = (Wet soil – Dry soil)/Wet soil, following Donagema et al. (2011).

Water table (WT) depth was directly measured in the floodable areas by digging shallow holes in the soil, when necessary. For the intermediate and dry plots, we also measured the slope variance, taking the WT depth of the nearby floodable area as a reference, and estimated the WT depth of the intermediate and the dry plots according to the variance detected along the soil slope.

### Variable selection

Overall, we measured 25 soil and moisture variables as environmental predictors of plant functional traits and forest trait composition. Of the plant traits measured, we chose to investigate aboveground biomass, community weighted specific leaf area, height, and basal area (AGB, SLA, height and BA). The selection of the most important environmental variables related to these traits was performed with generalized additive models (GAM) and generalized linear models (GLM) following analysis scripts in Neves et al. (2017). This technique uses (1) a forward selection method of environmental variable for redundancy analysis (RDA); (2) additional and progressive elimination of collinear variables based on their variance inflation factor (VIF) and ecological relevance, maintaining only those with VIF<4 (Quinn et al. 2002). VIF was calculated using the *R*^2^ value of the regression of one variable against all other explanatory variables. The GAM, GLM, NMDS, and ordination analyses were conducted in the R statistical environment (R Core Team 2018), using the packages “vegan” (Oksanen et al. 2018) and “recluster” (Dapporto et al. 2015).

### Structural equation modeling

Multiple analytical methods used in conjunction with one another to allow us to assess the effects of the most important environmental drivers on forest trait composition. Statistically significant environmental variables were forward-selected in GAM analysis for each response variable (AGB, SLA, BA and height). Following the variable selection through forward selection analysis (Blanchet et al. 2008), we constructed piecewise structural equation models (SEM) (Lefcheck 2016), and carried out independent effects analysis (Grace et al. 2012). SEMs allow hypothesis testing about the network of causal relationships between observables in a system and their relative strengths, without knowing ahead of time which variables will have positive, negative, direct, or indirect effects on those observables.

Piecewise SEMs were used to reveal relationships between predictor variables (soil properties) and response variables (plant traits) (Michaletz et al. 2018). The piecewise SEM analysis is performed with AIC model selection (Shipley 2013), and is better than traditional SEM analysis for small datasets, and those containing non-independent observations of multivariate non-normally distributed variables (Michaletz et al. 2018). These properties apply to our datasets for AGB, height and BA (Appendix S1. Fig. S4). We used “piecewiseSEM” (Lefcheck 2016) and “hier.part” (Walsh and Nally 2013) packages for R software environment (R Core Team 2018) to carry out the construction and analysis of the piecewise structural equation model.

## Results

### Measured trends in soil properties

All physical soil properties shifted substantially with water table depth, with the exception of clay content (Appendix 1. Fig. S1). We found that the proportion of coarse sand increased with water table depth, whereas fine sand and silt proportion decrease toward the drier end of the gradient, reducing the soil water retention capacity (Appendix 1. Fig. S1). Soils became impoverished in nutrients (Zn, Al, Na, Ca, Cu, SOM, CEC, SB, H+Al and ASI) with increasing water table depth (in drier environments). Exceptions to this trend were P, Mn, and B, which increased slightly with water table depth. All communities were found to have strongly acidic soils (pH<5), and there was a decreasing trend of soil acidity toward the drier communities (which had higher pH and lower H+Al) (Appendix 1. Fig. S2).

### Variable selection and structural equation modeling

GAM and SEM analysis (Table 2, Fig. 2 and Fig. 3) reveal that the functional composition of plots was primary affected by soil parameters related to water availability (WT depth and coarse sand), whereas soil nutrients (Fig. 2d), and Al, Na, and H+Al (Figs. 3 and 4) had a secondary role on the response variables. Interestingly, elements that are usually considered toxic (aluminum) or stressful for plants (sodium) were selected into the models (Table 2). Basal area, height, and AGB were negatively impacted by coarse sand soil content (Fig.2a, c and b). However, sodium saturation index (SSI) had a positive effect on AGB and height (Fig.2a, 2b, and 4). Potential acidity (H+Al) had a positive effect on the basal area (BA), despite its influence on the increase of Al soil availability, which negatively impacted BA (Figs. 2c and 3).

**Figure 2.**
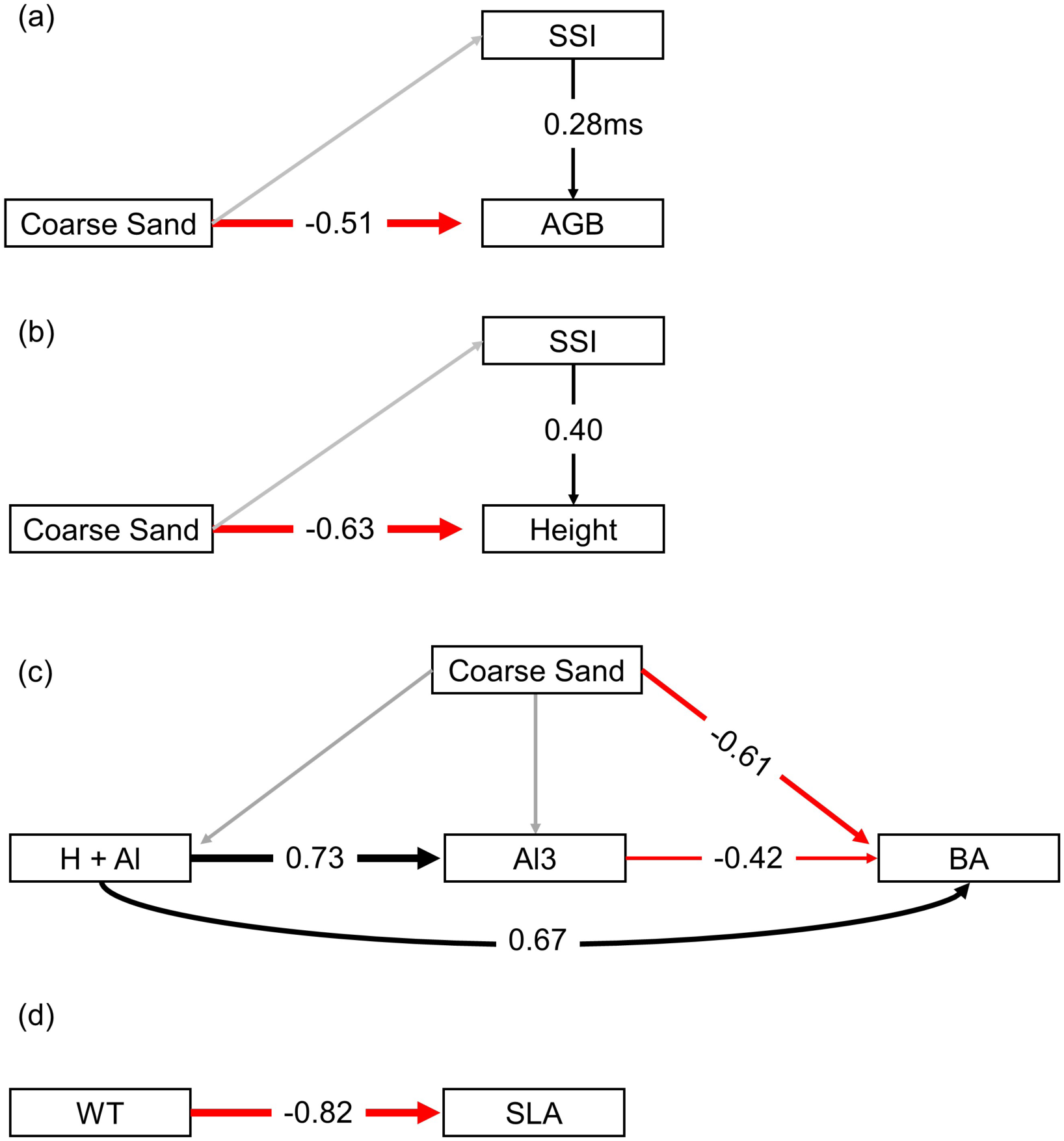
Simplified models obtained by piecewise structural equation models, using the forward selection analysis applied to all measured environmental variables. Shown are models for the most important drivers of (a) AGB, (b) height, (c) BA and (d) SLA from 42 *restinga* plant communities. Sodium’s effect on AGB was marginally significant (ms; *P* = 0.054). Water table depth (WT) had the dominant effect on SLA, and the remaining 24 environmental variables had low explanatory power with respect to SLA.

**Figure 3.**
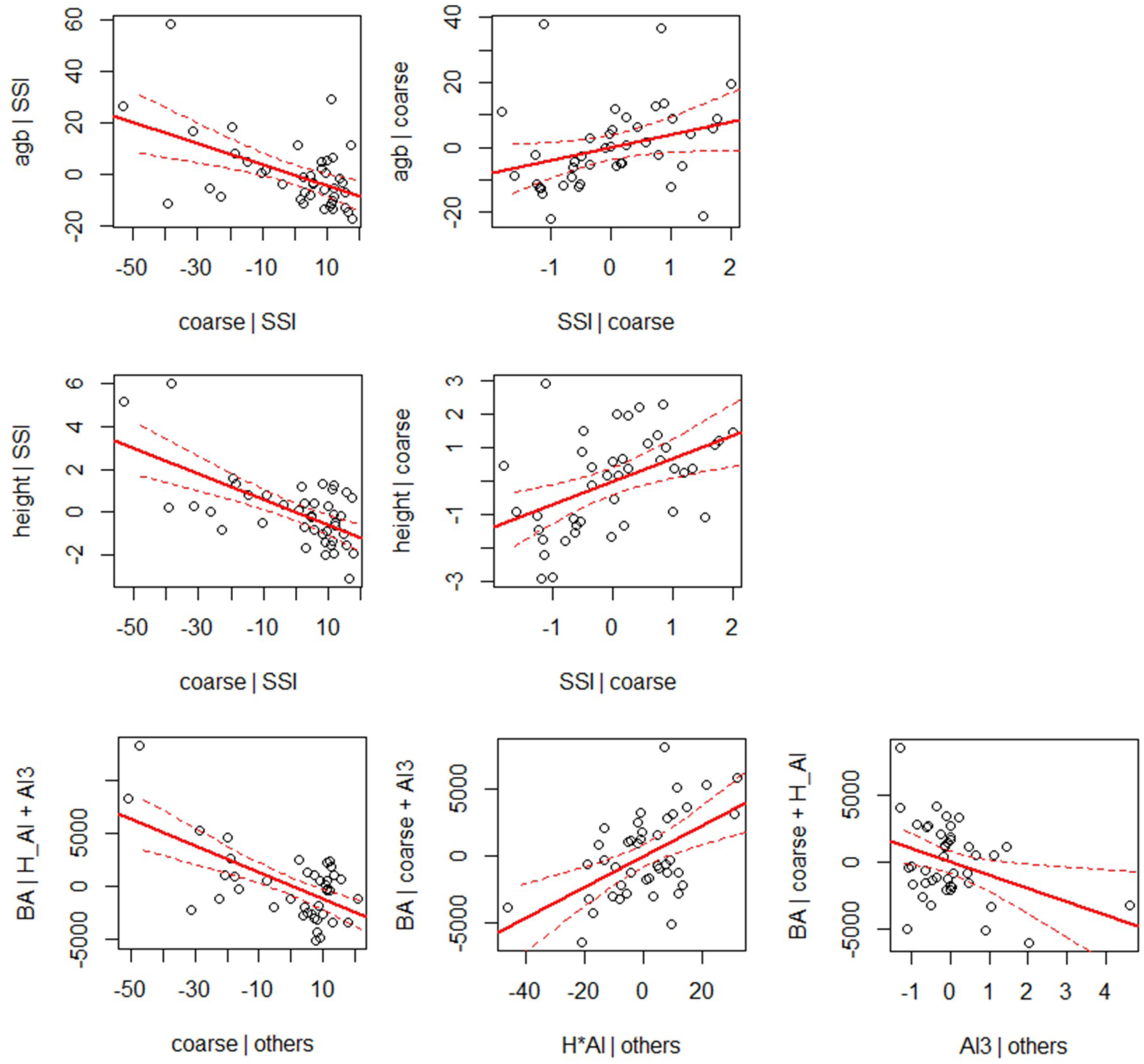
Drivers of aboveground biomass (AGB), basal area (BA), specific leaf area (SLA), and height in 42 *restinga* local plant communities. Relationships shown are the result of piecewise structural equation modeling. Partial correlations of direct effects are shown in (a), for example, there is a negative relationship between the effect of coarse sand (given SSI) on AGB (given SSI). The water table depth has a direct and strong effect on SLA (b), and water table depth was the only variable selected in the forward selection analysis.

**Figure 4.**
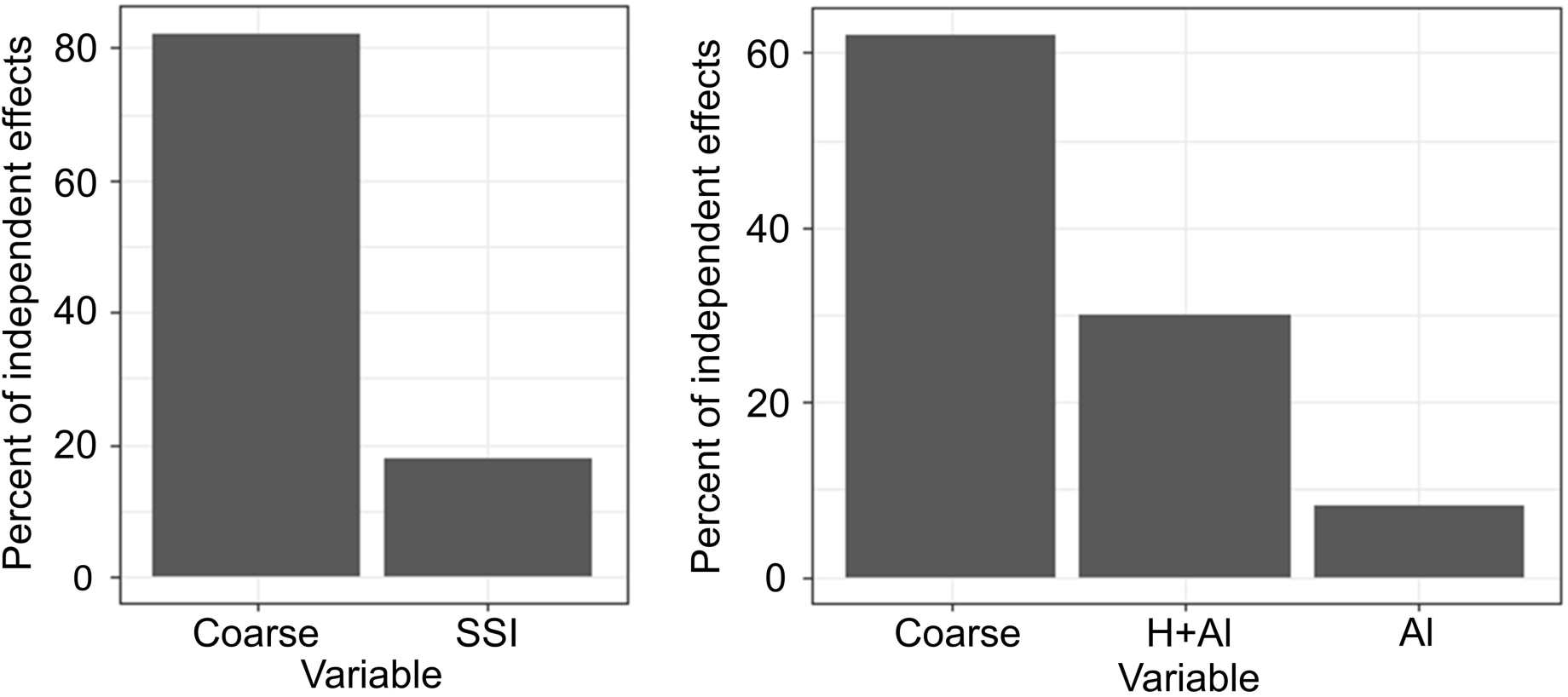
The separate independent effect of coarse sand, sodium saturation index (SSI), and aluminum (Al) on the aboveground biomass (AGB), basal area (BA), and plant height in 42 *restinga* plant communities. The separate independent effect is calculated by piecewise structural equation model selection via d-separation and Akaike’s information criterion (Lefcheck 2016, Shipley 2016). Positive numbers (black arrows) represent positive effects of one variable on another, and negative numbers (red arrows) represent negative effects. Larger magnitudes indicate larger effect sizes.

**Table 2.** Environmental variables related to aboveground biomass (AGB), height, specific leaf area (SLA), and basal area. Goodness-of-fit of predictor variables was assessed through adjusted coefficients of determination, the Akaike information criterion (AIC), *F*-test values and significance tests (*p* < 0.01 in all cases). Adj. *R*^2^ cum. = cumulative adjusted coefficient of correlation, represents total variance explained by each model.

| **AGB ~** | **adj. $R^2$ cum.** | **ΔAIC** | ***F*** | **Pr(>*F*)** |
| --- | --- | --- | --- | --- |
| Coarse sand | 0.182 | 216.99 | 10.13 | 0.006** |
| Sodium Saturation Index | 0.238 | 214.92 | 3.96 | 0.050* |
| <All variables> | 0.509 |  |  |  |
| **Height ~** |  |  |  |  |
| Coarse sand | 0.268 | 34.476 | 16.008 | 0.002** |
| Saturation Sodium Index | 0.410 | 26.332 | 10.665 | 0.004** |
| <All variables> | 0.596 |  |  |  |
| **SLA ~** |  |  |  |  |
| Water table depth (WT) | 0.640 | 321.84 | 73.999 | 0.002** |
| <All variables> | 0.681 |  |  |  |
| **Basal Area ~** |  |  |  |  |
| Coarse Sand | 0.249 | 679.83 | 14.644 | 0.002** |
| H + Al | 0.364 | 673.80 | 8.225 | 0.002** |
| Al3+ | 0.431 | 670.03 | 5.589 | 0.028* |
| <All variables> | 0.672 |  |  |  |

Increases in water table depth, coarse sand and aluminum soil concentration negatively impacted all the measured functional traits, whereas SSI had a positive effect on AGB and plant height. SLA was mainly affected by WT depth, which produced a high level of explained variation in SLA (64% of 68.1%), considering that 25 environmental variables were used in the global test of the forward selection step in the GAM analysis (Table 2). Overall, the models explained a high proportion of the variation in AGB (50.9%), BA (59.6%), and plant height (67.2%) (Table 2).

## Discussion

We found a few general trends defining strong edaphic gradients in *restinga* soils. Toward the dry environments, soils become sandier and more well-drained (Appendix S1. Fig. S1), and poorer in SOM, Zn, Al, Na, Ca and Cu, whereas P, Mn and B soil content increases slightly (Appendix S1. Fig. S2). Nutritional impoverishment toward the drier communities has been related to increased leaching of nutrients in well-drained soils of *restinga* plant communities (Lourenço, Jr. and Cuzzuol 2009, Magnago et al. 2010), as textural characteristics are highly associated with the retention of soil nutrients in *restinga* soils (Cooper et al. 2017). Furthermore, we measured base saturation (BS) lower than 10 (Appendix S1. Fig. S2), indicating low nutritional soil reserves, which may be even more impoverished in *restinga* soils that in similar tropical soils because of high rainfall and the sandy soil texture (Rodrigues et al. 2013).

We found that all soils were strongly acidic (Appendix S1. Fig. S2), with pH<5, and measured only slight changes of pH across the gradient. Below a pH of 5, severe depletion of the availability of essential nutrients as N, P, K, Ca and Mg has been observed (Lambers et al. 2008). However, as a higher concentration of hydrogen cations in the soil modestly increases the nutrients input by increasing weathering rates of soil organic matter (Lambers et al. 2008), floodable forest plant communities may take advantage of higher acidity in organic soils and the release of nitrogen compounds from the SOM. This may explain the positive effect of potential acidity (H+Al) on forest basal area (Fig. 2C and 3A). Moreover, flooding imposes anoxic conditions to the soil, which reduce the microbial decomposition activity and delay the nutrients releasing from SOM (Fridman 2005). Low rates of decomposition favors a more efficient absorption of nutrients by plant root systems, which may enhance the nutrient use in (Bonilha et al. 2012) and influence plant species distribution (de Almeida Jr et al. 2011).

We did not find evidence to support a primary role of SOM in plant community trait composition (Table 2). The most explanatory soil variables for AGB, SLA, basal area, and height were those related to the water table depth and soil drainage or coarseness (proportion coarse sand), whereas SOM was not selected as highly significant in the models we constructed. Compared to WT depth and soil drainage, SSI and aluminum had lower explanatory power and independent effects on trait composition (Table 2 and Fig. 3b). Thus, water availability-related soil parameters (WT depth and coarseness) seem to have a key role in the trait composition of such systems, while SSI and Al seem to exert a secondary role.

In a discussion about essential nutrients, Subbarao et al. (2003) present several arguments to include Na within the definition of functional nutrient, which is “an element that is essential for maximal biomass production or can reduce the critical level of an essential element by partially replacing it in an essential metabolic process.” In conditions of soil nutrients shortage or even in high Na soil content, Na can replace K, given their chemical similarity. Both elements compete for the same absorption sites in the plant root system and Na can assume some K-related metabolic functions, such as osmoticium for cell enlargement and accompanying cation for long-distance transport (Subbarao et al. 2003). Thus, the high Na soil content we measured (SSI>4.5%, and Na>90 g/kg; Appendix S1. Fig. S2), and its positive effect on AGB and height (Figs. 2a, 2b, and 3a) suggests that Na may be exerting a vital role in *restinga* plant nutrition, possibly supplementing the nutritional lack of K soil content (Appendix S1. Fig. S2). Moreover, as salinity was recently reported to distinguish *restinga* forests from others environmentally marginal habitats of Atlantic Forest (Neves et al. 2017), we may conclude that Na plays a key role in the system functioning, as has been observed in other topical forests (Kaspari et al. 2009).

The aluminum soil content in the study area (Fig. 3) had higher values (>6 cmol/kg) than those reported in earlier studies of *restinga* plant communities (Magnago et al. 2013, Rodrigues et al. 2013, Melo Júnior and Torres Boeger 2015). Al solubility and availability is known to be related to the increase of soil acidity (Lambers et al. 2008); and we found that Al availability was higher toward the wet end of the flooding gradient (Appendix S1. Fig. S2), where soils also showed strong acidity (pH = 3.6). Moreover, the potential acidity (H+Al) exerted a positive effect on Al soil content, which in turn produced a negative effect on BA (Fig. 2b). This finding is supported by similar studies, in which the combination of lower pH (<5.5) and higher Al soil concentration results in toxic conditions for many plant tissues, constraining growth, and the uptake of several essential nutrients (US Environmental Protection Agency 2014, Bojórquez-Quintal et al. 2017).

It has been demonstrated that the application of Al can stimulate the uptake of N, P and K in acidic soils (Osaki et al. 1997), enhancing the growth of some native plant species that have adapted to acidic conditions through exclusion or accumulation of Al (Watanabe and Osaki 2006), which was reported for some species found in *restinga* forests, including *Tapirira guianensis* (Britez et al. 2002a) and *Faramea marginata* (Britez et al. 2002b). Other benefits of Al in Al-adapted plants include increased defense against pathogens, alleviation of abiotic stress, and increased metabolism and antioxidant activity (Bojórquez-Quintal et al. 2017).

The accumulation of aluminum in leaves and seeds, and the positive effect of the soil available aluminum in plant survival have been reported in some native plants from Brazilian *Cerrado* ecosystem (Haridasan 2008), which shares several plant species with *restinga* forests, such as those we found in our study area: *Tapirira guianensis*, *Calophyllum brasiliense*, *Protium heptaphyllum*, *Myrcia rostrata*, *Xylopia sericeae*, *Alchornea triplinervia*, *Pseudobombax grandiflora*, *Emmotum nitens*, *Pera glabrata*, *Amaioua guianensis*, and *Cecropia pachystachya* (Lourenço et al., *in prep.*). The similarity of both systems with regard to the high aluminum soil content and the co-occurrence of several plant species reinforce our findings of the importance of aluminum in the functioning of *restinga* forests (Figure 4).

Despite of the several environmental variables used in this study and the high level of retained explanation for all response variables (Table 1, adj. R^2^ cum. ranging from 51% to 68%), a considerable amount of variation remained unexplained (32% to 49%). These findings suggest the existence of other factors beyond the physicochemical soil properties influencing forest physiognomy and functional composition, highlighting the importance of the biotic interactions, such as competition (Garbin et al. 2016) and facilitation (Garbin et al. 2014), and climatic drivers, whose investigation could further broaden our understanding of the drivers of forest composition and function in the threatened coastal Atlantic Forest.

## Supplementary Material

Additional Supporting Information may be found in the online version of this article: **Appendix S1. Additional analyses of soil properties and plant functional traits.**

## Acknowledgments

This work was conducted at the University of Arizona and JLJ was financed in part by the Coordenação de Aperfeiçoamento de Pessoal de Nível Superior - Brasil (CAPES) - Finance Code 001 - PDSE program grant n° 88881.131961/2016-01. Funding for EAN came from the USDA Forest Service and the University of Arizona through the Bridging Biodiversity and Conservation Program.

## Supporting Information

### Appendix S1

**Table S1.**
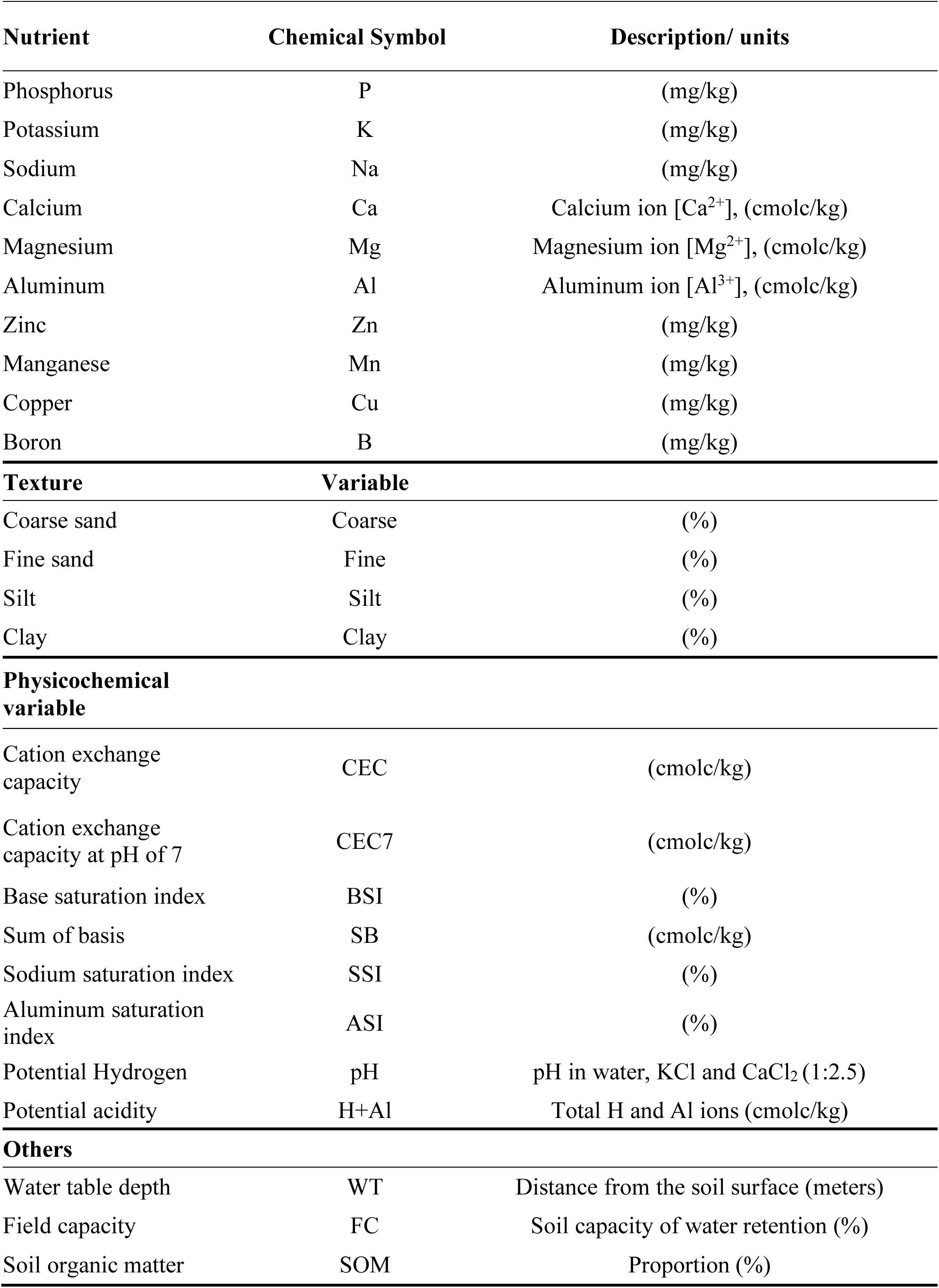
Environmental variables used to the forward selection analysis, including nutrients, texture, and physicochemical soil parameters, with a brief description and units.

| Nutrient | Chemical Symbol | Description/ units |
| --- | --- | --- |
| Phosphorus | P | (mg/kg) |
| Potassium | K | (mg/kg) |
| Sodium | Na | (mg/kg) |
| Calcium | Ca | Calcium ion [Ca<sup>2+</sup>], (cmolc/kg) |
| Magnesium | Mg | Magnesium ion [Mg<sup>2+</sup>], (cmolc/kg) |
| Aluminum | Al | Aluminum ion [Al<sup>3+</sup>], (cmolc/kg) |
| Zinc | Zn | (mg/kg) |
| Manganese | Mn | (mg/kg) |
| Copper | Cu | (mg/kg) |
| Boron | B | (mg/kg) |
| Texture | Variable |  |
| Coarse sand | Coarse | (%) |
| Fine sand | Fine | (%) |
| Silt | Silt | (%) |
| Clay | Clay | (%) |
| Physicochemical variable |  |  |
| Cation exchange capacity | CEC | (cmolc/kg) |
| Cation exchange capacity at pH of 7 | CEC7 | (cmolc/kg) |
| Base saturation index | BSI | (%) |
| Sum of basis | SB | (cmolc/kg) |
| Sodium saturation index | SSI | (%) |
| Aluminum saturation index | ASI | (%) |
| Potential Hydrogen | pH | pH in water, KCl and CaCl<sub>2</sub> (1:2.5) |
| Potential acidity | H+Al | Total H and Al ions (cmolc/kg) |
| Others |  |  |
| Water table depth | WT | Distance from the soil surface (meters) |
| Field capacity | FC | Soil capacity of water retention (%) |
| Soil organic matter | SOM | Proportion (%) |

**Figure S1.**
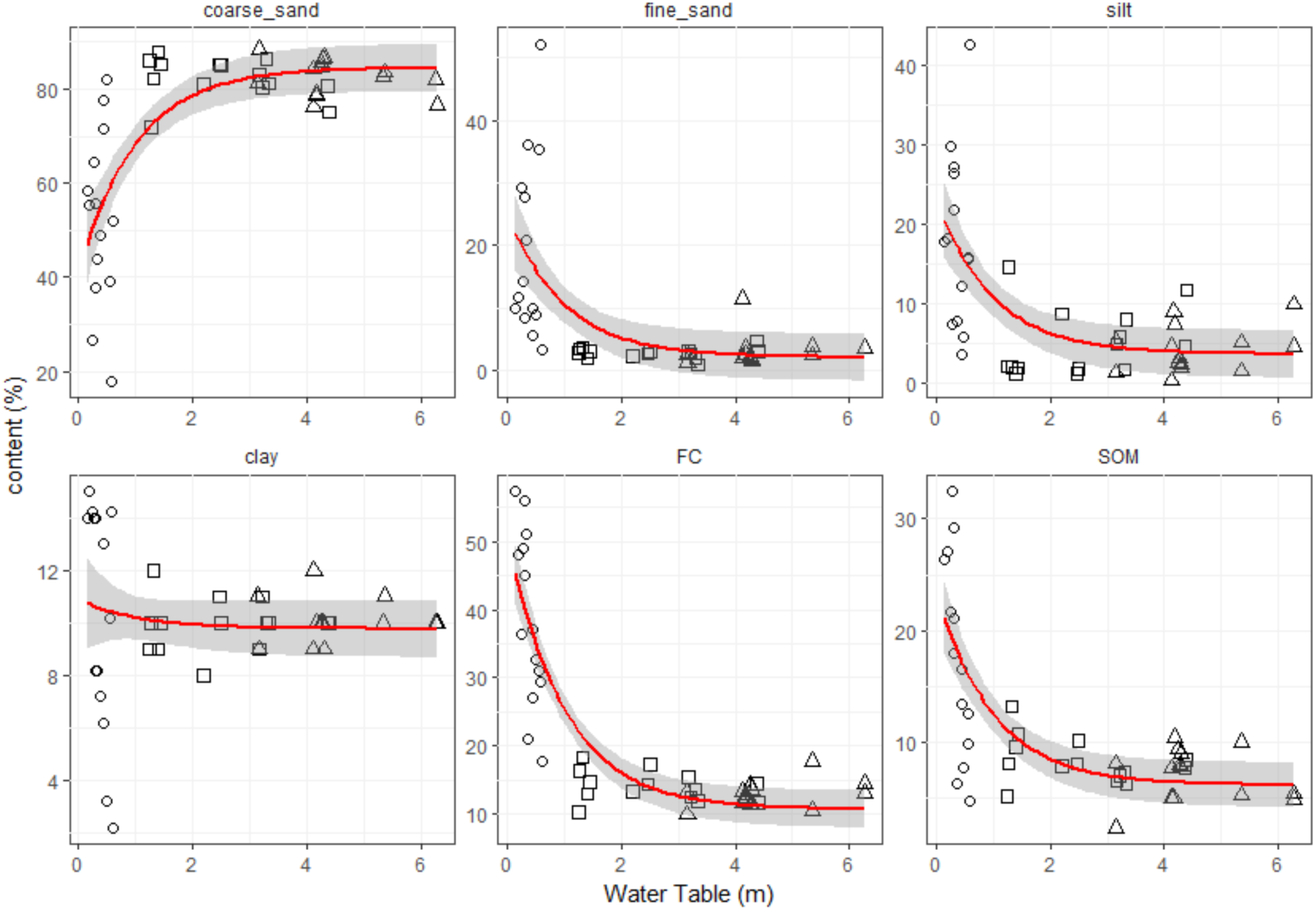
Soil coarseness, field capacity (FC) and soil organic matter (SOM) variation across water table gradient. Communities from floodable (○), intermediate (□), and dry (Δ) sites.

**Figure S2.**
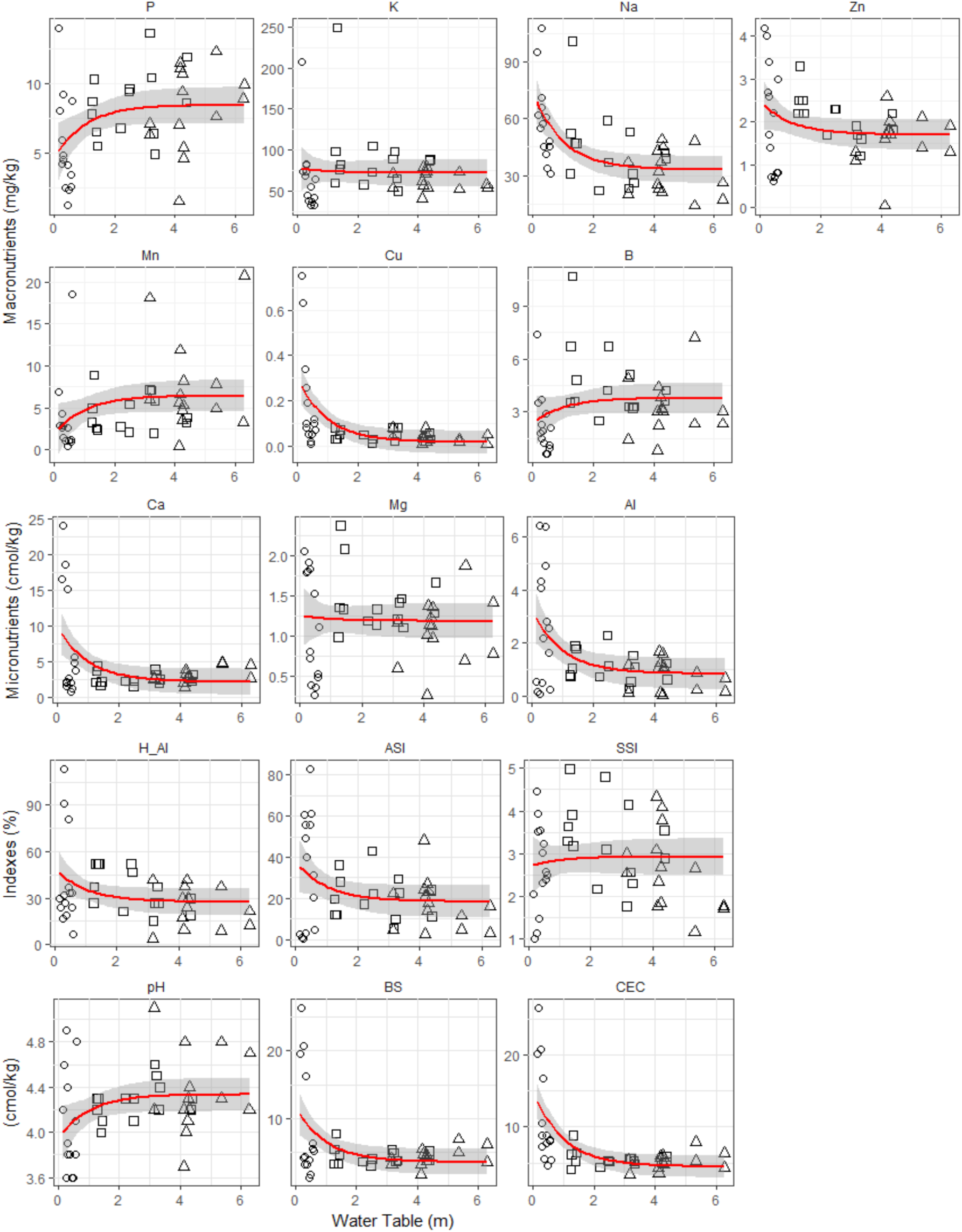
Variation of soil chemical properties across the water table gradient, showing macronutrients; micronutrients; acidity (pH), cation exchange capacity (CEC), base saturation (BS), acidity related to hydrogen and aluminum (H+Al), aluminum saturation index (m), sodium saturation index (SSI). Plots from floodable (○), intermediate (□), and dry (Δ) sites.

**Figure S3.**
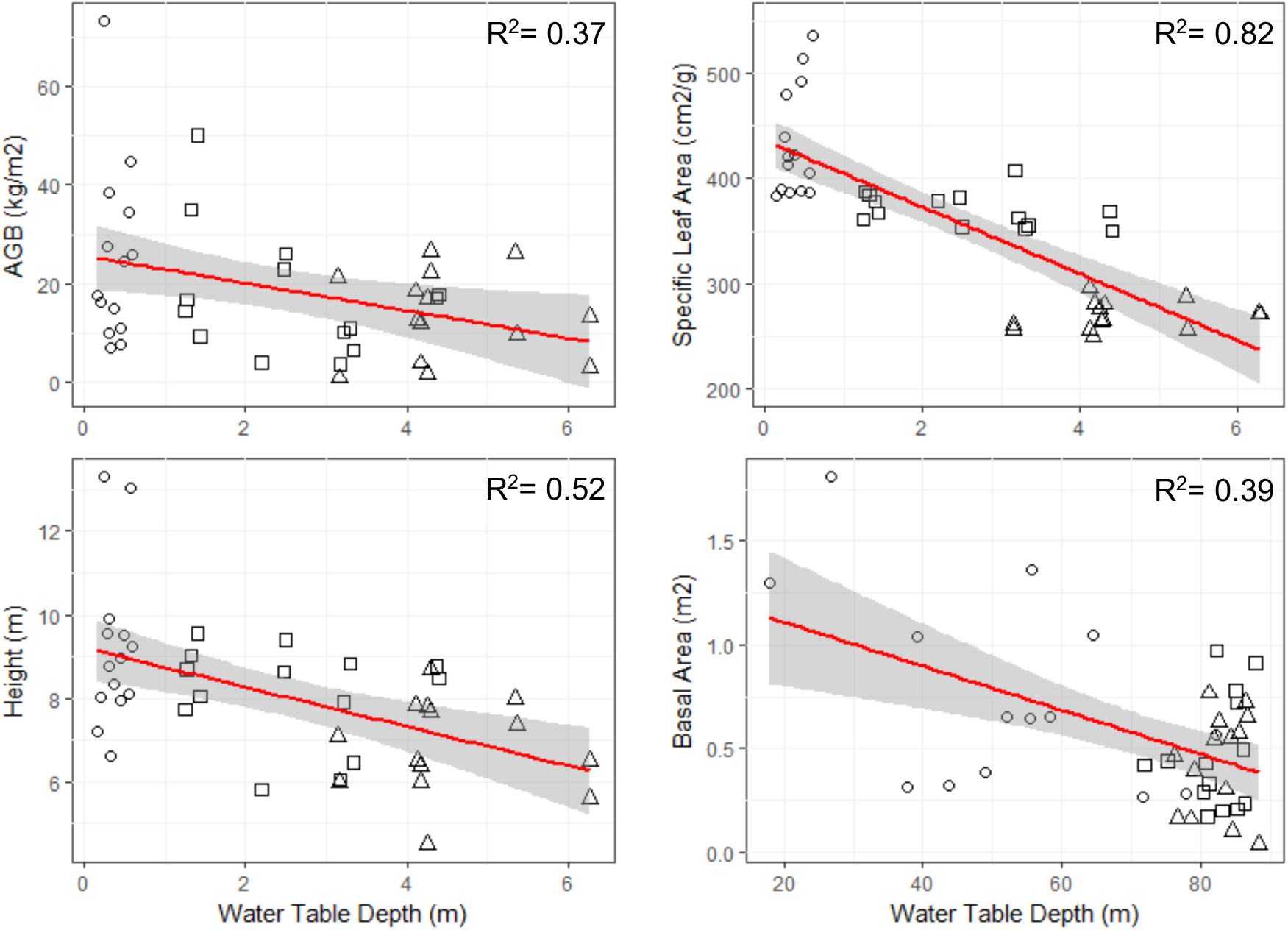
Aboveground biomass (AGB), community weighted specific leaf area (SLA), height, and basal area (BA) distributions along the water table depth gradient. All correlations were statistically significant (*P-*value < 0.05). Each dot represents one of the 42 plots from floodable (○), intermediate (□), and dry (Δ) sites.

**Figure S4.**
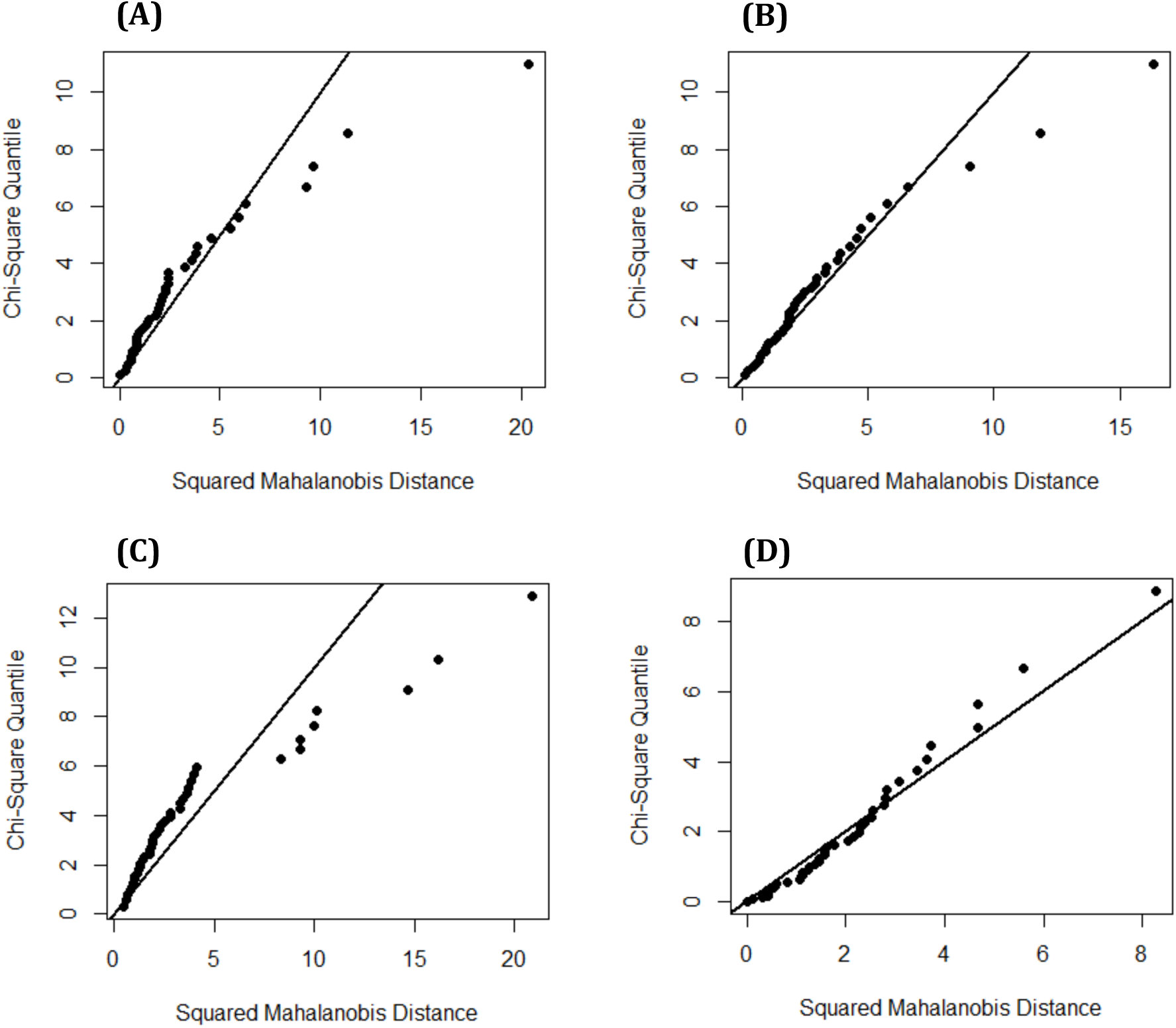
Chi-Square Q-Q plot of the variables from each equation model: **(A)** AGB ∼ coarse sand + SSI; **(B)** height ∼ coarse sand + SSI; **(C)** BA ∼ coarse sand + (H+Al) + Al3; and **(D)** SLA ∼ water table. All the models except **(D)** show departures from multivariate normality.

Multivariate non-normality is assessed by Mardia’s test for variables, with a null hypothesis of multivariate normality and a test statistic that will have an approximate Chi-squared distribution. Deviations from the Chi-squared distribution indicate lack of multivariate normality. For model **(A)** (*P_skew_* = 0.004, *P_kurtosi_*_s_ = 0.86); model **(B)** (*P_skew_* = 1.54, *P_kurtosis_* = 0.02); and model **(C)** (*P_skew_* = 3.73 x 10^-12^, *P_kurtosis_* = 3.21 x 10^-9^). The variables from the model **(D)** showed multivariate normality (*P_skew_* = 0.35, *P_kurtosis_* = 0.28). Statistical tests were conducted using the package “MVN” (Korkmaz et al. 2014) in the statistical software R (R Core Team 2018).

